## Supplementary for "Evolutionary constraints on the complexity of genetic regulatory networks allow predictions of the total number of genetic interactions"

### Supplementary information

**Authors:** Adrian I. Campos-González[1,2] and Julio A. Freyre-González[1,*]

[1]Regulatory Systems Biology Research Group, Laboratory of Systems and Synthetic Biology, and [2]Undergraduate Program in Genomic Sciences, Center for Genomics Sciences, Universidad Nacional Autónoma De México, Av. Universidad s/n, Col. Chamilpa, 62210, Cuernavaca, Morelos, México.

**Supplementary note 1: The May-Wigner stability theorem**

In the early 1970s, Gardner and Ashby empirically found that the stability of randomly connected large systems depends on their connectance[1]. They explored the system stability by modeling a set of nonlinear first-order differential equations whose coefficients, representing the interaction strengths among variables, were randomly obtained from a Gaussian distribution having zero mean and variance $\alpha^2$. The change of the system state $\mathbf{x}(t)$ (where $\mathbf{x}(t) = (x_1(t), x_2(t), \ldots, x_n(t))$) in time can be represented by the equation $d\mathbf{x}/dt = \mathbf{A}\mathbf{x}$, where $\mathbf{A}$ is the matrix of interaction strengths. The percentage of non-zero entries in this matrix was defined as the percentage of connectedness (connectance). Connectance is then analogous to density in graph theory as both quantify the fraction of existing interactions relative to the total possible.

Connectance (and consequently also density) has an important role in complexity theory as it quantifies the complexity of a system[1,2]. Robert M. May extended Gardner and Ashby's work to conclude that the stability of randomly connected systems depends on the number of variables ($n$), connectance ($C$), and interaction strength dispersion ($\alpha^2$)[2]. The May-Wigner stability theorem says that randomly connected large systems are stable if $nC < 1/\alpha^2$. Unstructured and structured systems have been shown to be bound by this theorem. An early theoretical work in structured model networks has suggested that such structures promote stability[3], and this was also observed for hierarchical networks under certain conditions[4]. However, a later theoretical study concluded that hierarchical and modular networks are less stable than random networks[5] and other showed that increasing modularity or the number of hierarchical layers tends to increase the probability of instability[6]. It has been theoretically suggested that optimizing multiple structural and dynamical constraints such as minimizing complexity and path length while increase robustness to dynamical perturbations will evolve modular scale-free networks[7].

**Supplementary note 2:** ***Fitting xmin parameter for MLE of power-law distribution.***

The maximum likelihood estimate MLE for a power-law probability distribution has been derived[8,9] and is implemented in a python package called *powerlaw*[10]. Because of the parametrization of a power-law distribution, very small values' probability would tend towards infinity. The valid areas for which a power law would suffice a probability distribution are defined by a parameter called *xmin*. Although theoretically a valid approach for estimating the parameters for fitting power-law distributions, we identified an undesired behavior arising from using this *data trimming* parameter when comparing Erdos-Renyi (ER) and biological networks P(k). To explain the phenomena, and why setting the *xmin* parameter to 1 was proposed as a solution, we will discuss an example. Let us take *Escherichia coli* 2017 GRN including its RNA-mediated interactions (accession number: 511145_v2017_sRDB16_dsRNA), and generate an ER equivalent network having the same number of nodes and density. Notably ER graphs are constructed so that their node degree distribution follows a Poisson distribution. A histogram of the degrees of both networks would look like this:

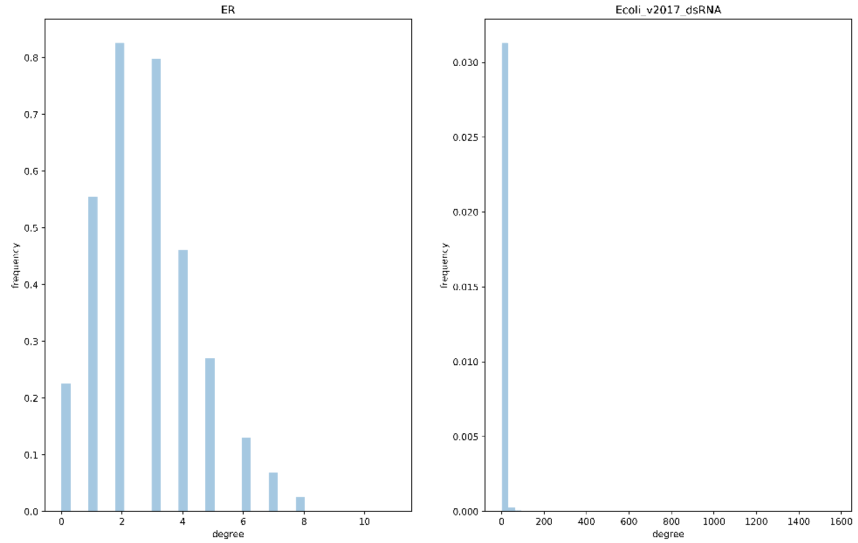


From this figure, we can already notice differences between the node degree distributions of these networks. First, while ER network seems normally distributed, *E. coli* GRN seems to follow a decaying trend and second, the biological network obviously follows a long-tailed distribution while the Poisson derived one does not.

If we were to fit a power-law distribution to the ER P(k) allowing for *xmin* to be a free parameter, most of the data of the ER network gets trimmed out. Importantly the trimmed data no longer seems to have a good fit to a poisson distribution (see below right panel).

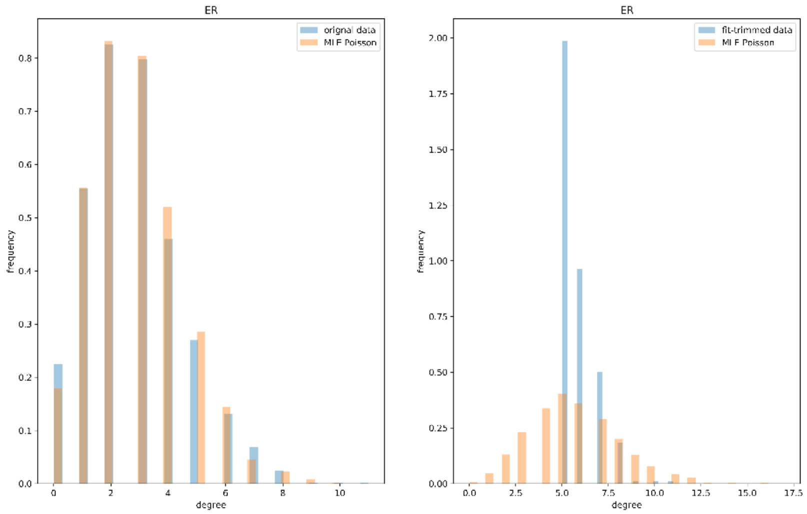


Allowing for a free *xmin* impedes us from understanding the true fit of the data to long-tailed distributions. Furthermore, a measure of the amount of information being ignored is not directly available from the estimates. Although in this example the *xmin* for ER and Biological networks are fairly close (5 and 3 respectively), their effects on trimming the data are far from being equal:

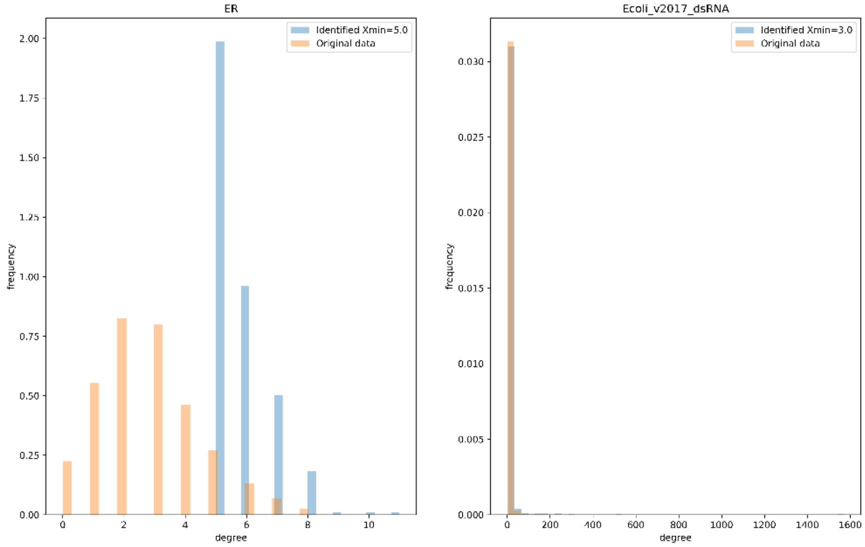


This result motivated us to repeat the analyses using a fixed *xmin* parameter of one, ensuring the use of all data available both for biological and theoretical networks.

### Supplementary figures

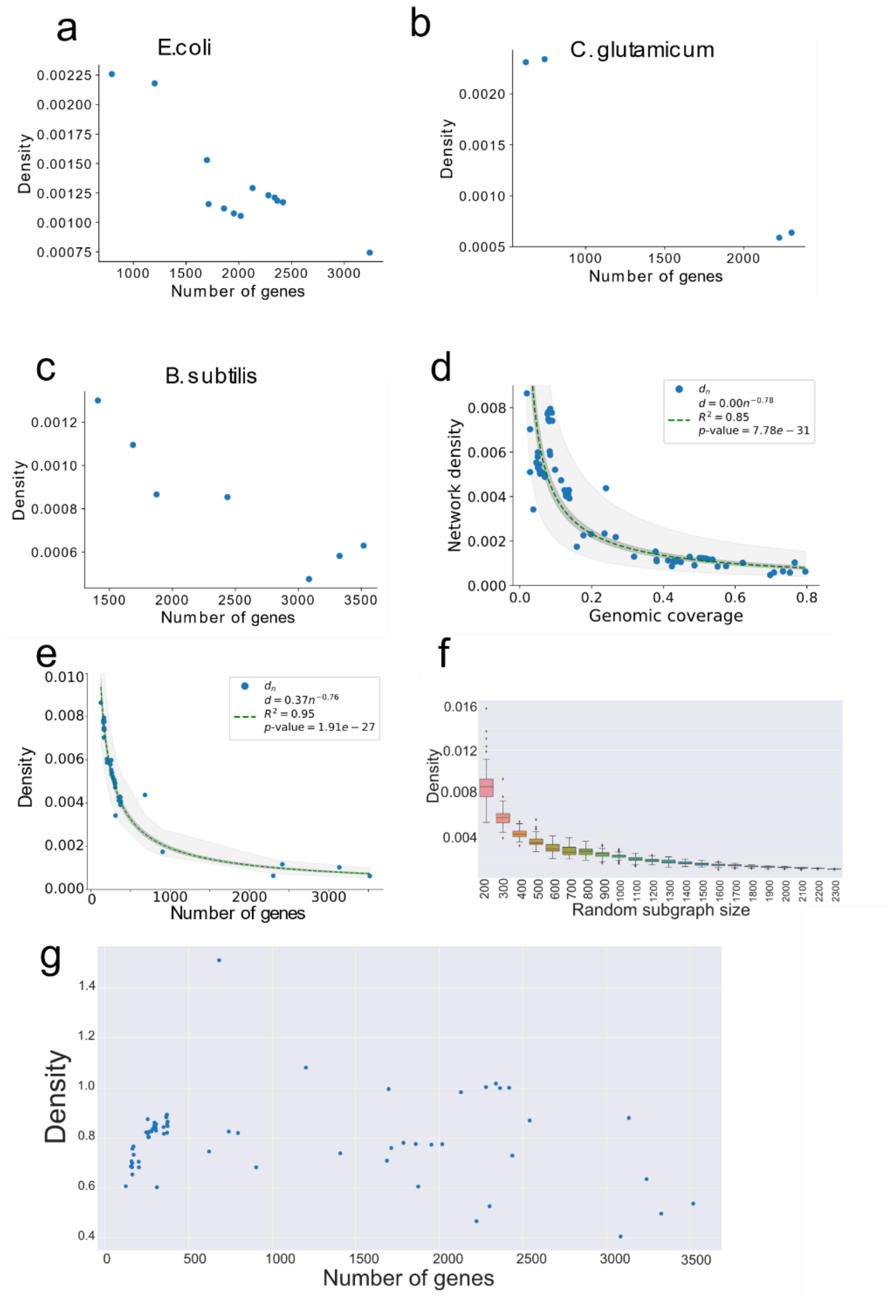


**Supplementary figure 1. Abasy GRNs present a trend towards a low density.**

a) Network density decreases with network completeness in historical reconstructions of *E. coli*. The same results are recapitulated in reconstructions of *C. glutamicum* (b) and *B. subtilis* (c). d) The use of genomic coverage as a completeness proxy does not affect the trend observed in network densities. e) The same density trend (as Fig. 1a) is observed when using a set of non-redundant GRNs. f) Random (snowball) sampling of *E. coli* 2013 network generates the same pattern as observed in (a), explaining the variability in networks with lower genomic coverage. g) Invariance observed in density when normalizing by randomly expected number of nodes in a GRN.

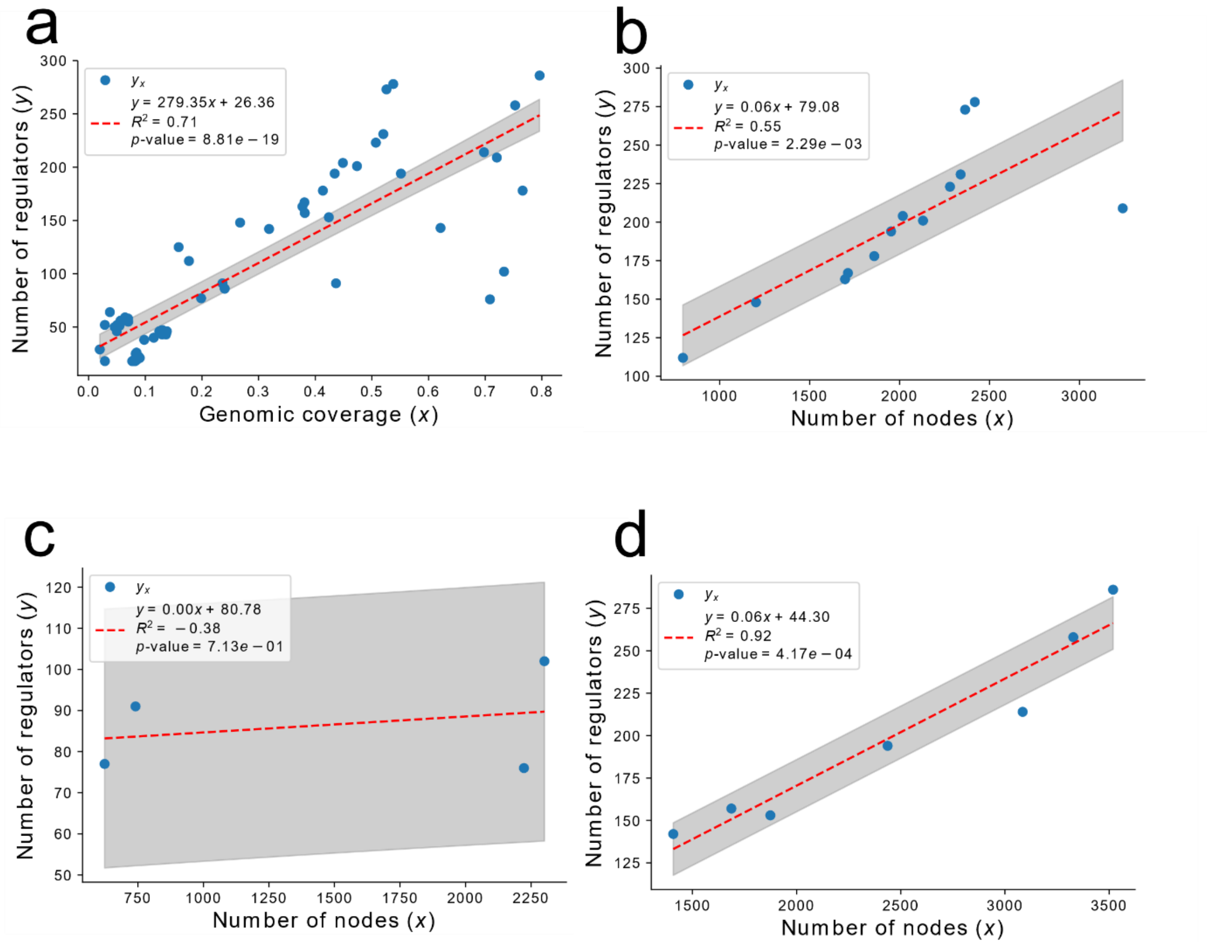


**Supplementary figure 2. Relationship between number of regulators and genes.**

The observed trend between completeness and number of regulators was recapitulated when using genomic coverage to assess completeness (a). The significant trend was also present in a set of historical reconstructions of *E. coli* (b), *C. glutamicum* (c) and independent *B. subtilis* (d) GRNs. We acknowledge a lack of power (data) for the historical reconstructions of *C. glutamicum* but included this result for reproducibility and openness.

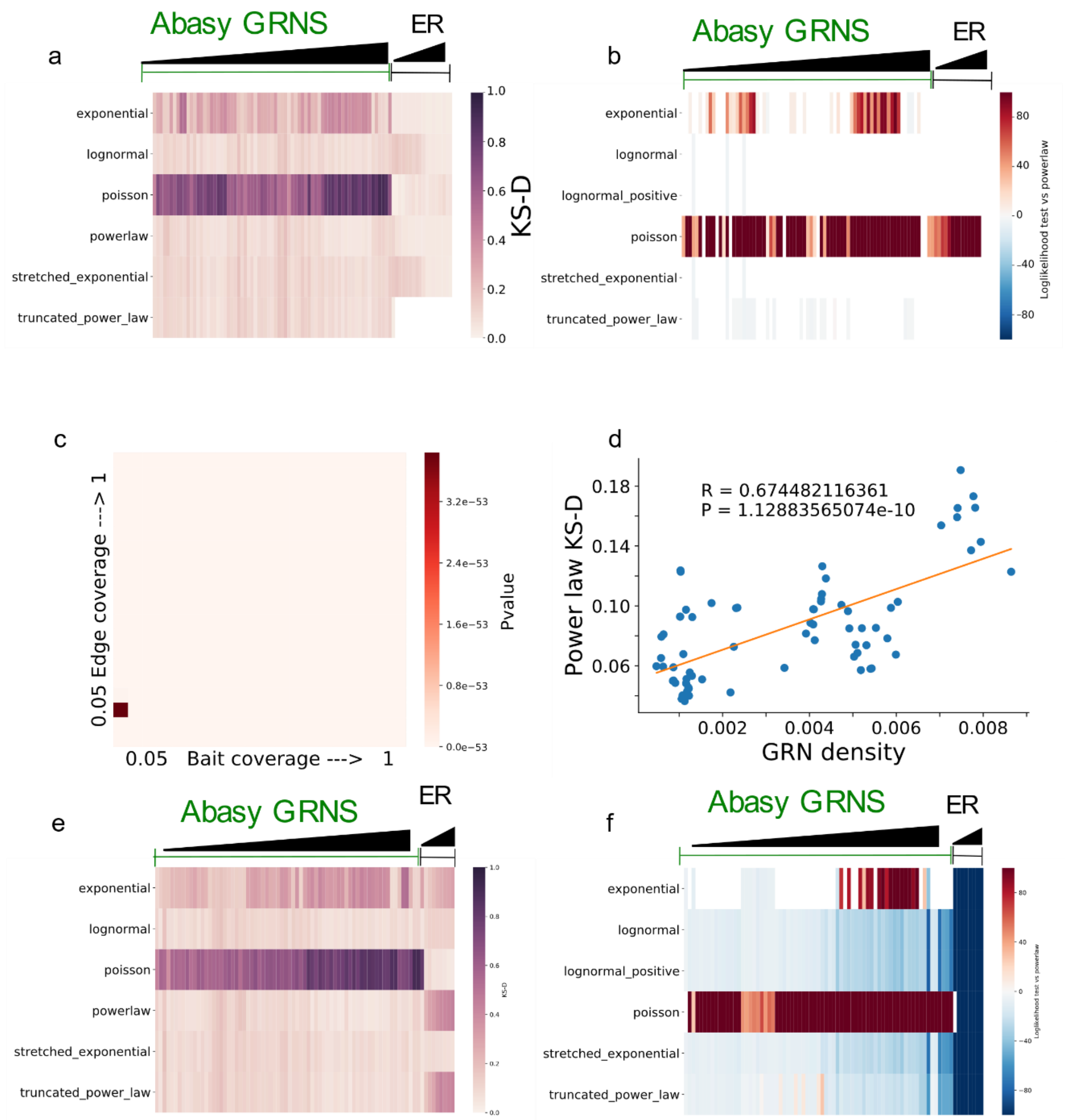


**Supplementary figure 3. GRN P(k) distribution follows a power-law distribution not likely arising from an artifice of sampling.**

a,b) Effect of the trimming parameter xmin when assessing goodness of fit of ER graphs to a set of probability distributions. These results correspond to the same approach as in **Figure 2**, but allowing xmin parameter to vary, thus not fitting all the connectivity data. Note that this causes **all** ER networks to have a good fit to and preference for a power law even when they are derived from a Poisson (see ER graphs and compare with **Figure 2**. c) P-values of the KS_D differences reported on **Figure 2e** note that all values are very significant. d) A negative correlation between goodness of fit to a power law and graph density was observed for the GRNs in Abasy. e,f) The results presented in Figure2 are recapitulated when using another sampling scheme for the ER networks: snowball sampling (**see methodology**).

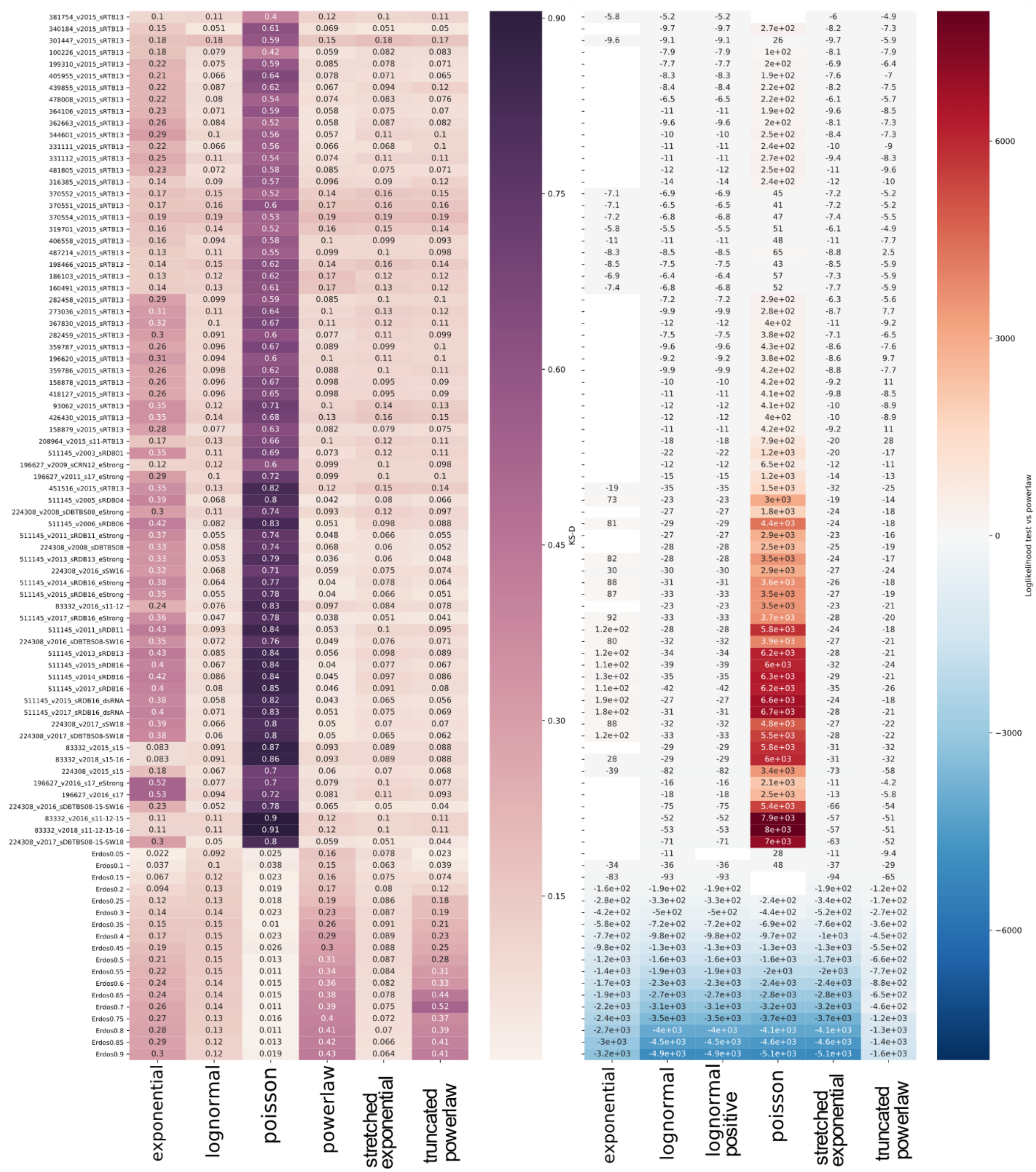


**Supplementary figure 4. Annotated version of Figure 2 from main manuscript.**

A detailed view with annotation scores is provided here.

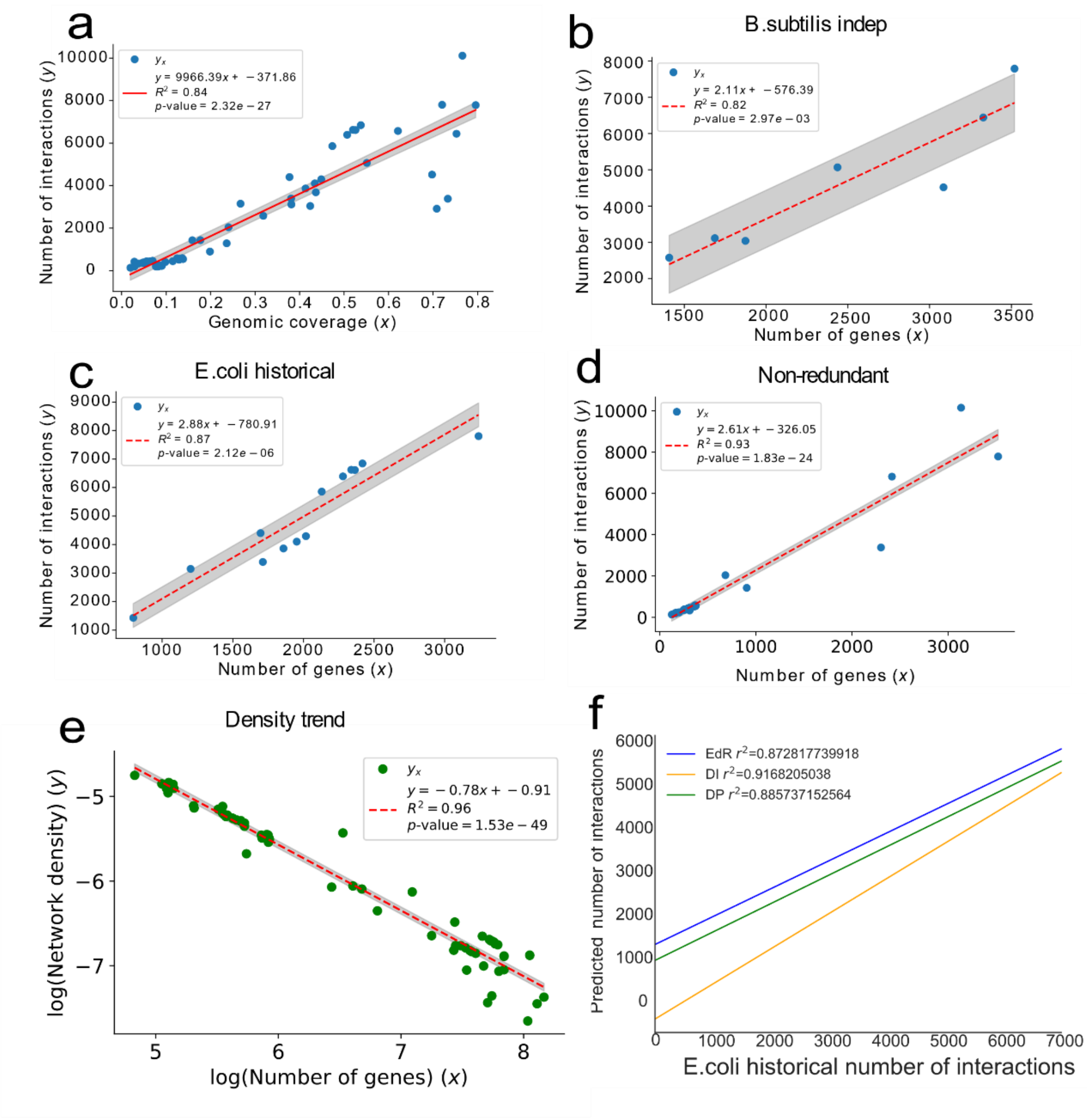


**Supplementary figure 5. Model generation and profiling.**

a) Relationship between genomic coverage and number of interactions. b and c) show the relationship between nodes and edges in historical reconstructions of *E. coli* and *B. subtilis*, respectively. d) Relationship between number of genes and number of interactions in the set of non-redundant networks. e) Density proportionality model parametrization. We model density as an exponential decay (by fitting a linear regression to the log transformed values) to predict number of nodes (**Fig 4c**). f) Comparison of the performance of the three models predicting *E. coli* historical reconstruction number of edges. Briefly, the three models were used to predict the number of interactions of the different *E. coli* GRNs and their accuracy was measured by the residual squared error ($R^2$)

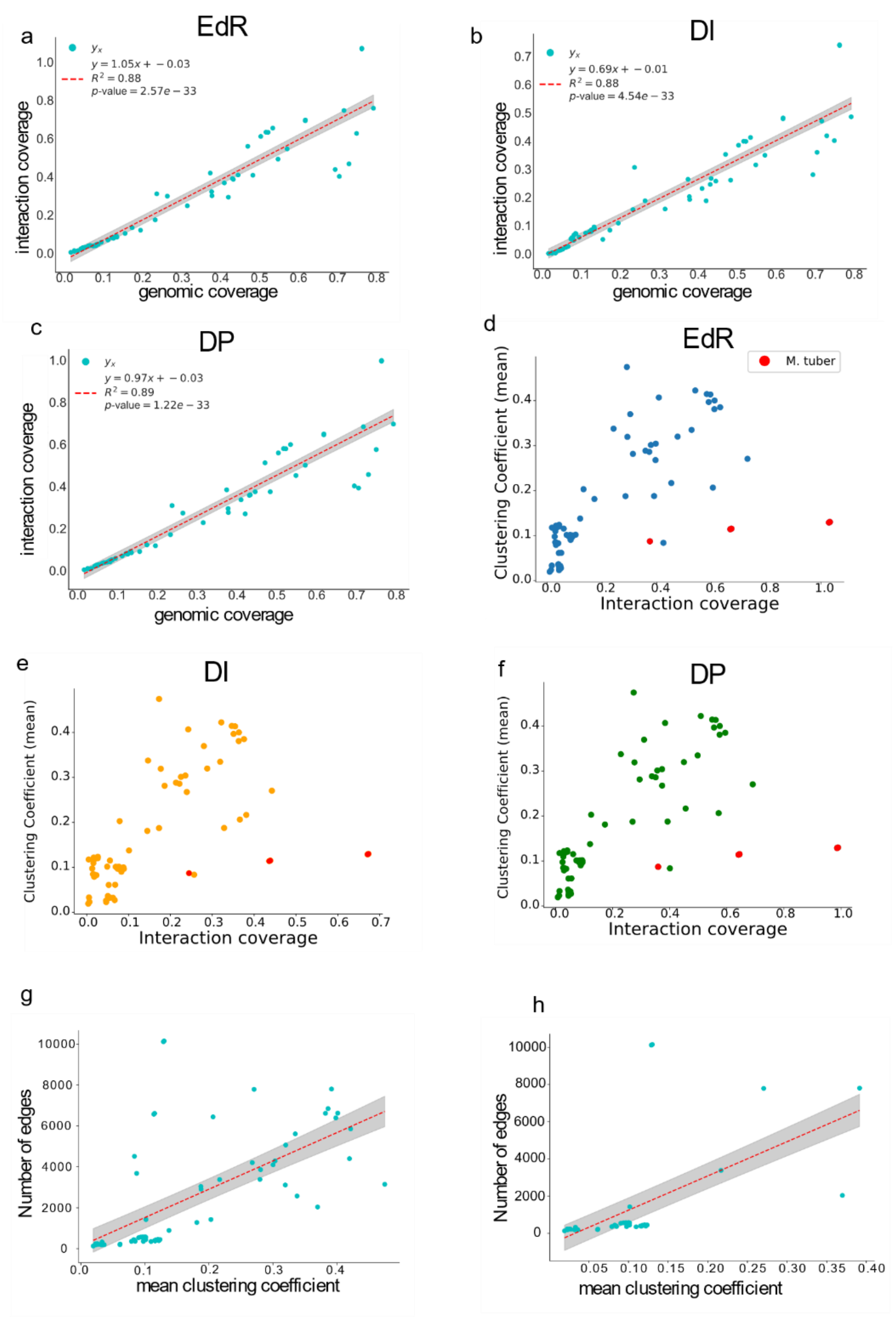


**Supplementary figure 6. Relationship between interaction coverage and clustering coefficient.**
a-c) A high correlation between the interaction coverage and genomic coverage using the edge regress, density invariant and density proportionality models respectively. d-f) Relationship between interaction coverage and mean network clustering coefficient using as estimator the models assuming EdR, density DI and DP models, respectively. g) Relationship between number of edges and clustering coefficients in Abasy GRNS. h) Same as in g but with the non-redundant networks.

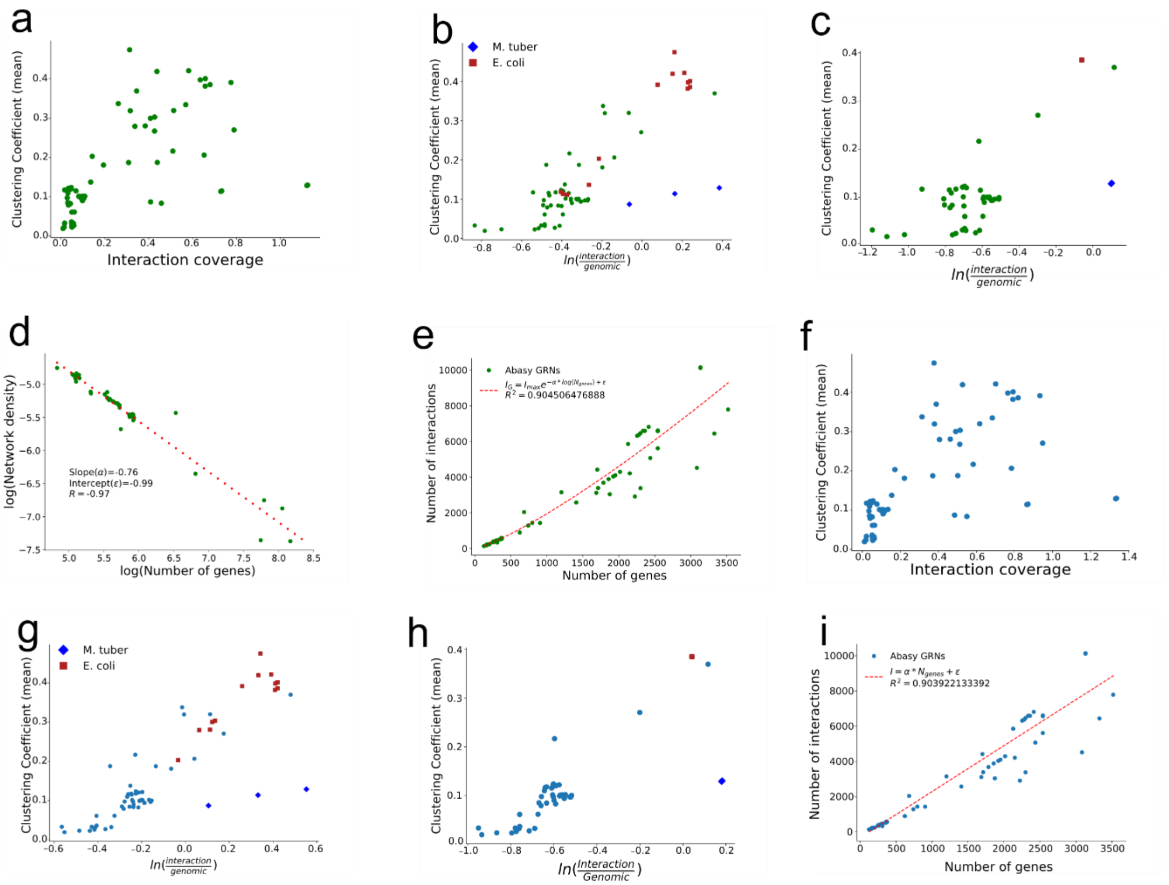


**Supplementary figure 7. Completeness estimation based on non-redundant networks.**

Model parameters were computed again using only a set of non-redundant, most complete networks (**see methods**). All panels are based on estimates using this set of networks. a-e) Model characteristics of density proportionality model. Panel b and c depict the same relationship, but on all Abasy GRNs (b) or only on the subset of non-redundant networks used to fit the models (c). d) Depicts the new parameters for modelling DP as an exponential decay to predict (e) number of interactions. f-i) Edge regress (EdR) model results when using only a set of non-redundant networks to parametrize the model. Note that overall results are similar to the ones presented in the main text using all networks to fit the models.
